# Amyloid accumulation drives proteome-wide alterations in mouse models of Alzheimer’s disease like pathology

**DOI:** 10.1101/150623

**Authors:** Jeffrey N. Savas, Yi-Zhi Wang, Laura A. DeNardo, Salvador Martinez-Bartolome, Daniel B. McClatchy, Timothy J. Hark, Natalie F. Shanks, Kira A. Cozzolino, Mathieu Lavallée-Adam, Samuel N. Smukowski, Sung Kyu Park, Jeffery W. Kelly, Edward H. Koo, Terunaga Nakagawa, Eliezer Masliah, Anirvan Ghosh, John R. Yates

**Author notes:** Correspondence: (J.N.S.) or (J.R.Y.). Lead contact: (J.R.Y.). Current Addresses: Department of Molecular Physiology and Biophysics, Vanderbilt University, School of Medicine, Nashville, TN 37232, USA. Howard Hughes Medical Institute and Department of Biology, Stanford University, Stanford, CA 94305, USA. E-Scape Bio. South San Francisco, CA 94080, USA.

## Abstract

Amyloid beta (Aβ) peptides impair multiple cellular pathways in the brain and play a causative role in Alzheimer’s disease (AD) pathology, but how the brain proteome is remodeled during this process is unknown. To identify new protein networks associated with AD-like pathology, we performed global quantitative proteomic analysis in three mouse models at pre- and post-symptomatic ages. Our analysis revealed a robust and consistent increase in Apolipoprotein E (ApoE) levels in nearly all transgenic brain regions with increased Aβ levels. Taken together with prior findings on ApoE driving Aβ accumulation, this analysis points to a pathological dysregulation of the ApoE-Aβ axis. We also found dysregulation of protein networks involved in excitatory synaptic transmission consistent with AD pathophysiology. Targeted analysis of the AMPA receptor complex revealed a specific loss of TARPγ-2, a key AMPA receptor trafficking protein. Expression of TARPγ-2 *in vivo* in hAPP transgenic mice led to a restoration of AMPA currents. This database of proteome alterations represents a unique resource for the identification of protein alterations responsible for AD.

**Highlights:**

- Proteomic analysis of mouse brains with AD-like pathology reveals stark remodeling
- Proteomic evidence points to a dysregulation of ApoE levels associated with Aβ clearance rather than production
- Co-expression analysis found distinctly impaired synapse and mitochondria modules
- In-depth analyses of AMPAR complex points to loss of TARPγ-2, which may compromise synapses in AD

**eTOC Blurb:** Proteome-wide profiling of brain tissue from three mouse models of AD-like pathology reveals Aβ, brain region, and age dependent alterations of protein levels. This resource provides a new global protein expression atlas for the Alzheimer’s disease research community.

## Introduction

Alzheimer’s disease (AD) is a progressive brain disorder that is the leading cause of dementia in adults, resulting in impaired memory and cognition. Pathological hallmarks of AD include the presence of amyloid plaques, neurofibrillary tangles, astrogliosis, and changes in vasculature in the brain, which ultimately culminate in neurodegeneration and premature death. Abundant evidence shows that at the biochemical, molecular, and cellular levels that AD brain pathology is highly complex and represents feedback and feed forward responses in multiple cell types (De Strooper and Karran, 2016).

There has been a plethora of potential mechanisms proposed to explain AD pathology, although a unifying and universally accepted description has yet to be achieved. There is a near unanimous agreement that amyloid beta (Aβ) peptides generated by proteolytic processing of the amyloid precursor protein (APP) represent a key toxic species; however, a precise mechanism for their damaging activity remains highly controversial. Historically, the “amyloid cascade hypothesis” has provided a useful theoretical framework for the basis of AD (Selkoe and Hardy, 2016), yet current evidence suggests a more complicated pathological etiology (Herrup, 2015). One of the major challenges impeding our understanding of AD pathology is that many molecular perturbations vary based on the stage of disease progression, cellular context, and brain region. Furthermore, even individuals with causative mutations in APP or Presenilin-1 (PS1) may take multiple decades to develop AD, which suggests that long incubation periods and / or aging is required before neuronal homeostasis is irreversibly lost. Several molecular pathways including impaired synaptic transmission, hampered protein degradation dynamics, defective mitochondrial function, and inflammatory responses all play important roles in AD pathogenesis (Akiyama et al., 2000; Lin and Beal, 2006; Mori et al., 1987; Sheng et al., 2012).

Dynamic changes in protein abundances represent essential cellular responses needed to cope with cellular insults and stress. However, broad toxicity, such as the accumulation of Aβ peptides, is likely to affect many proteins and pathways. We hypothesized this must ultimately result in proteome remodeling (Powers et al., 2009). For example, proteins physically associated with Aβ plaques are known to accumulate during disease progression while other proteins such as those localized to synapses are believed to have reduced levels due to the loss of synaptic contacts that occurs during AD pathology (Lassmann et al., 1992). The situation is further complicated by the numerous cell types in the brain, their interactions, and the potential mechanisms responsible for the discrete spreading of pathology. To understand the effects of Aβ peptides and signaling caused by amyloidogenic proteolytic processing of APP on the mammalian brain proteome, we first performed global quantitative proteomic analysis of frontal cortex, hippocampal, and cerebellar extracts from transgenic mice expressing mutated hAPP or hAPP / PS1 (Borchelt et al., 1996; Mucke et al., 2000). While these widely used models have provided breakthrough insight into AD pathology, they are potentially limited by the over-expression of APP fragments outside Aβ in non-relevant cells in the brain (Saito et al., 2014). To address this potential limitation, we have also performed proteomic analysis of BRI-Aβ42 mouse brain extracts that only over-express Aβ42 without additional APP fragments (McGowan et al., 2005). Furthermore, to temporally resolve changes in protein abundance, we performed our analysis at both pre-symptomatic and post-symptomatic time points to find unknown, early, and potentially reversible pathway alterations.

Here, we present an interactive AD proteomic resource consisting of 18,882 quantified proteins mapping to 10,288 genes from mouse model brain region extracts expressing hAPP, hAPP and PS1, or BRI-Aβ42. Through these efforts, we identified and confirmed many proteins never before reported altered in AD. To delineate proximal from down-stream effects of Aβ on the brain proteome we first determined Aβ levels and amyloid plaque loads and performed proteomic analysis at pre- and post-symptomatic time points from the same animals. Interestingly, we observed an age dependent increase in proteome remodeling and brain region specific patterns of altered protein levels. We identified many proteins that are genetically linked to sporadic AD, such as ApoE, as significantly altered in many of our datasets. ApoE has an important role in regulating Aβ metabolism and is co-deposited in senile plaques in patients with AD (Liu et al., 2013; Namba et al., 1991). ApoE also appears to play a role in Aβ aggregation since removal of the *APOE* gene eliminated fibril Aβ deposition in amyloid mouse models (Bales et al., 1997). However, whether ApoE levels are altered in AD patients remains an important and unclear issue (Gupta et al., 2011; Hesse et al., 2000; Liu et al., 2013). Interestingly we found ApoE to be significantly increased in all three models and only in those brain regions harboring increased levels of Aβ. Overall, we identified a small collection of proteins significantly altered in the hippocampus or frontal cortex in multiple models. Co-expression analysis revealed two distinct modules which are involved with synaptic transmission in the hippocampus and two modules which are related to mitochondria in the frontal cortex. Biochemical purification of AMPARs, which represented core module proteins in the hippocampus from the hAPP model and human AD brains, showed a specific loss of TARPγ2 (also known as CACNG2 or stargazin). In order to assess if the proteomic data can lead to important conclusions we tested and found that correcting TARPγ2 levels could restore glutamatergic synaptic transmission *in vivo*. Altogether, our proteomic results provide a global resource of altered protein levels in multiple mouse models of amyloid toxicity, which reinforces the complex pathology associated with Aβ accumulation and AD pathology.

## Results

### Remodeled proteomes in the brains of hAPP and hAPP / PS1 transgenic AD mouse models

To determine how Aβ peptide toxicity affects the brain proteome and to examine AD-like pathology *in vivo*, we have developed a quantitative proteomic strategy using nitrogen-15 (^15^N) labeling of whole mice (Wu et al., 2004). Efficient labeling of the mouse brain proteome was accomplished by feeding non-transgenic mice a ^15^N spirulina diet from three to seven months (mo) of age (McClatchy et al., 2007). The ^15^N “heavy” labeled mice were used as an internal standard for global protein quantitation of two different transgenic AD mouse models (Tg-AD) relative to non-transgenic (Non-Tg) litter mates which both remained unlabeled and thus “light”. The mixed “light” and “heavy” proteins were digested to peptides, separated using multi-dimensional chromatography, and analyzed by tandem mass spectrometry (MS) (Figure 1A). We compare the frontal cortex (FC), hippocampus (HIP), and cerebellum (CB) between age-matched Tg-AD or non-Tg litter mates using the same whole brain “heavy” reference proteome and we performed our analysis at pre- (3 mo) and post-symptomatic (12 mo) time points to identify early and also age dependent changes in protein abundance associated with AD-like pathology. In total, our MS analysis provided relative quantitation for > 3,800 different proteins in each dataset of three or more biological replicates per group (Figure 1B).

**Figure 1.**
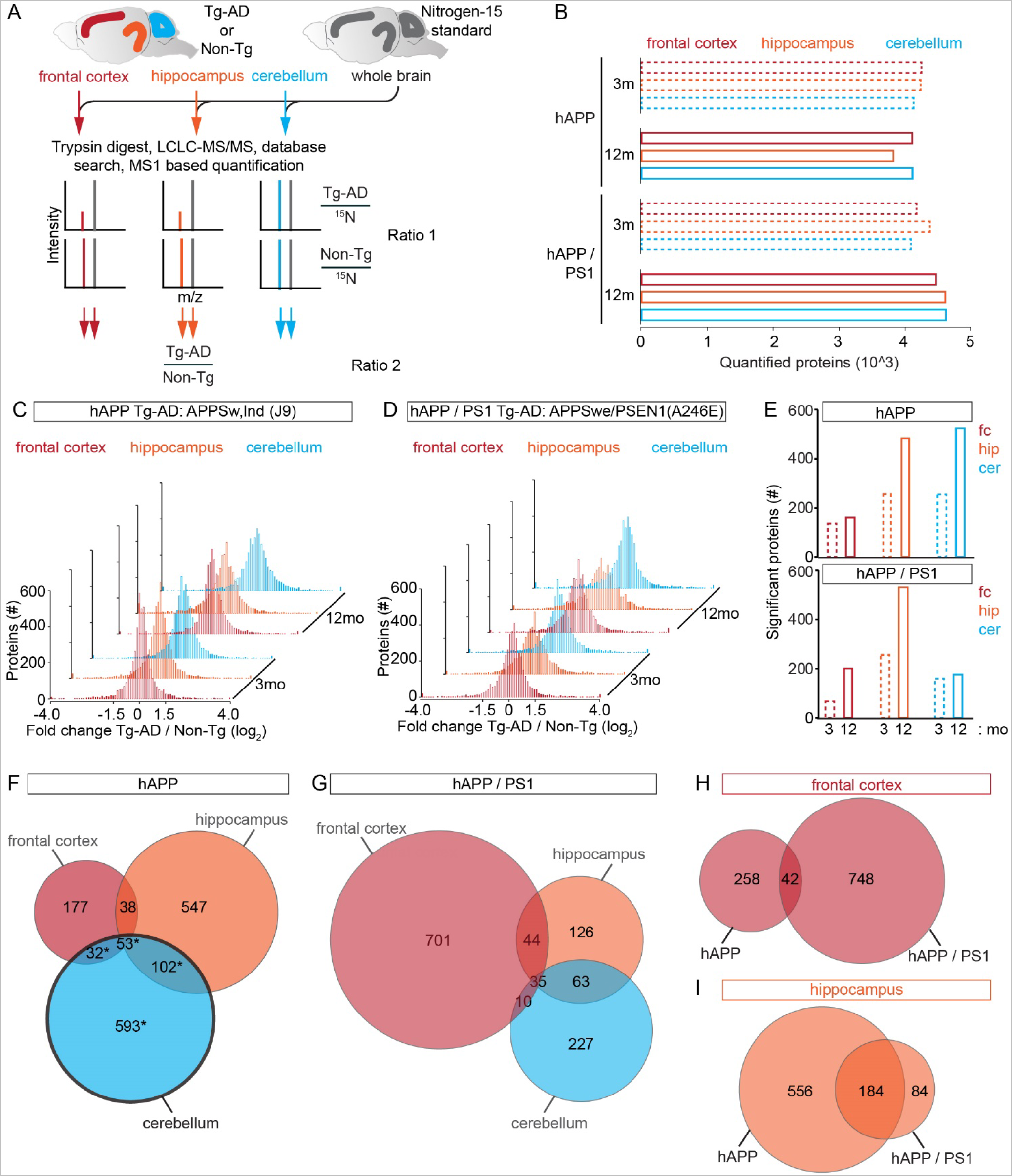
(See also Figures S1 and S6). Quantitative Proteomic Analysis of Three Brain Regions from Multiple Transgenic AD Mouse Models at Pre- and Post-symptomatic Time Points **(A)** Experimental design and analysis work flow. Frontal cortex, hippocampus, and cerebellum were dissected from 3 and 12 mo old Tg-AD or non-Tg litter mates and mixed 1:1 with ^15^N whole brain extracts. Proteins were digested into peptides and analyzed by shotgun proteomics and bioinformatics. The ratio of ^14^N to ^15^N is calculated and normalized to controls. **(B)** Summary of the number of quantified proteins in our 12 datasets. We quantified more than 3,800 proteins in each analysis. **(C)** Proteome remodeling of multiple brain regions in APP_swe/ind_ Tg-AD model mice at 3 and 12 mo. Solid bars represent those proteins with a > 50% change in ratio 2. **(D)** Proteome remodeling of multiple brain regions in APP_swe_ / PSEN1(A246E) model mice at 3 and 12 mo. Solid bars represent those proteins with a > 2-fold change in ratio 2. **(E)** Summary bar graphs showing an age dependent increase in the total number of significantly altered proteins (FDR-adjusted *p* value < 0.05) for each of the indicated datasets. **(F-G)** Venn diagrams indicating the number of proteins with significantly altered levels for the indicated brain regions and model, considering both time points. hAPP, black trace indicates cerebellum which lacks increased Aβ levels. **(H)** FC summary for both models, **(I)** HIP summary for both models.

By analyzing regional brain proteomes of two different Tg-AD models with different degrees of pathology we were able to identify new AD pathological mechanisms as well as confirming mechanisms that were previously reported. To identify proximal Aβ dependent protein perturbations, we first analyzed hAPP mice over expressing APP_swe/ind_ at moderate levels. These hAPP mice have been reported to have an onset of hampered synaptic transmission at the age of two to four months, but they do not develop predominant plaques before 10 mo of age (Hsia et al., 1999; Mucke et al., 2000). In addition, we performed quantitative proteomic analysis of a double Tg-AD mouse model over expressing both mutated hAPP and presenilin 1, specifically APP_swe_ / PSEN1(A246E), which we refer to as hAPP / PS1 (Borchelt et al., 1996). hAPP / PS1 mice have been reported to have plaques, dystrophic neurites, and behavioral deficits by 9 mo of age (Wang et al., 2003). To document the pathological state of our hAPP mice at 3 mo we performed Aβ 1-42 ELISA and found a significant, ~50%, increase in the level of Aβ 1-42 in the FC and HIP but not the CB (Figure S1A). At 12 mo Aβ 1-42 levels in the FC and HIP were even more significantly increased compared to the non-Tg littermates, while the cerebellum did not possess significantly increased Aβ 1-42 levels. The absence of a significant increase in the Aβ 1-42 levels in the CB of the hAPP model at both time points provides a useful negative control for proteins that are sensitive to APP fragments outside Aβ. Aβ 1-42 ELISA showed a large and significant increase in the Aβ 1-42 level in all three brain regions at both 3 and 12 mo in hAPP / PS1 (Figure S1B). Next, we analyzed the amyloid plaque load in hippocampal sections by Congo red and Thioflavin S staining of both Tg-AD models. This analysis confirmed previous reports that at 3 mo hAPP mice lack visible plaques, the number of plaques increased with age in both models, and the overall plaque load even at 12 mo is moderate for both models (Figure S1C-P). Based on these results we consider the HIP and FC datasets from the 3 mo hAPP model as predominantly pre-symptomatic, since we observed significantly increased Aβ 1-42 levels without detecting any visible amyloid plaques. Altogether the HIP and FC datasets from the 3 mo hAPP model best represent the earliest stage of pathology and proximal Aβ substrates.

Globally, our proteome-wide analysis of both Tg-AD models revealed a substantial imbalance in protein levels, with about 5% of the quantified proteins having a change of > 50% at 3 mo and an even larger percentage altered > 50% at 12 mo in 11 out of 12 of our comparisons (Figures 1C-D). To obtain accurate measurements of protein abundance, we used an average of at least 30 different peptides measurements per protein in each dataset (Figure S1Q). The total number of significantly altered proteins was increased from 3 mo to 12 mo in all brain regions in both models (hAPP: FC 3 mo = 138 and 12 mo = 162, HIP 3 mo = 256 and 12 mo = 484, CER 3 mo = 254 and 12 mo = 526, hAPP / PS1: FC 3 mo = 68 and 12 mo = 200, HIP 3 mo = 257 and 12 mo = 533, CER 3 mo = 159 and 12 mo = 176), and overall the HIP to be the brain region possessing the greatest number of significantly altered proteins (FDR-adjusted *p* value < 0.05) (Figure 1E). We compared the levels of all the significantly altered proteins in the same Tg-AD model and brain region between the 3 and 12 mo datasets. Interestingly, in the FC of hAPP but not hAPP / PS1 mice, we found that the difference between the average protein fold change at 3 to that at 12 mo was significantly increased (-0.0621 + 0.389 versus 0.214 + 0.320; mean + STD; *p* value = 1.34 E-8), consistent with an overall accumulation of protein in the context of more severe pathology (Figure S1R-S). For the HIP, we found a significant increase between 3 and 12 mo in hAPP / PS1 (-0.266 + 0.619 versus -0.155 + 0.604 mean + STD; *p* value = 0.0240). For the CB we found a significant increase in the hAPP / PS1 model, (-0.00866 + 0.554 versus 0.209 + 0.489; mean + STD; *p* value = 3.34 E-4). Next, we determined the number of proteins which were found significantly altered in each brain region and identified those proteins altered in multiple brain regions. Overall each brain region in both models had hundreds of proteins with significantly altered levels (Figures 1F-G and Table S1-2). We used bioinformatic analysis to investigate which cellular compartments the altered proteins localize and found that they reside in wide range of cellular compartments (Figure S1T). We then asked if any proteins were found significantly altered in the FC of both models and thus independently confirmed, indeed, 42 proteins were significantly changed. (Figure 1H and Table S3). Similarly, for the HIP we found 184 proteins significantly altered in both models (Figures 1I and Table S3). To determine how many of our altered proteins represented novel findings, we performed a comprehensive literature search and determined that 17 FC and 92 HIP proteins have never before been reported as being specifically altered in AD or AD models (Table S3). Altogether, these results reveal dramatic brain region specific proteome remodeling in Tg-AD mouse brains and provide confirmation of altered protein levels for those proteins found in multiple analyses.

We used hierarchical clustering based on the centroid method, to compare regional expression profiles for proteins involved with synapses, mitochondrion, or inflammation all of which have all been implicated in AD (De Strooper and Karran, 2016), we found robust brain region specific expression patterns (Figures S2A-C). Principal component analysis supported this finding and showed that for the FC the 3 and 12 mo proteomes of both models are most similar to each other, for the CB both time points for hAPP or hAPP / PS1 were most similar, and the HIP datasets were least similar between the models and time points (Figures S2D-F). The observation that the HIP datasets were the least similar between the two models could presumably be due to the effects of dissimilar Aβ levels (Figure S1A-B). These results demonstrate that our proteomic analysis strategy has sufficient resolution to identify brain region specific alterations potentially relevant to AD pathology.

### Connecting protein abundance changes to genes that predispose for AD

Next, we extended the potential connection between genes that have been linked to sporadic AD predisposition and changes in protein abundance in our proteomic datasets (Bertram et al., 2007). Our screen identified twenty-three significantly altered proteins which when mutated have been genetically linked to late onset AD (Figure 2A). Of these genes ApoE and Glutamic-Oxaloacetic Transaminase 1 (Got1) were found altered in six of our datasets, Glutathione S-Transferase M1 (Gstm1) was found in 5, and transferrin (TF) in 4. To further refine this observation, we extracted the model, time point, and brain region where these alterations were observed. Interestingly, we found that ApoE was significantly altered only in those datasets with significantly increased Aβ levels and other proteins such Bridging Integrator 1 (Bin1) and Clusterin (Clu) which have been tightly associated with predisposing for sporadic AD were only found altered in the more severe hAPP / PS1 model (Figure 2B). These findings show that a large number of genes genetically linked to AD also have altered protein products in mouse model brains of AD-like pathology.

**Figure 2.**
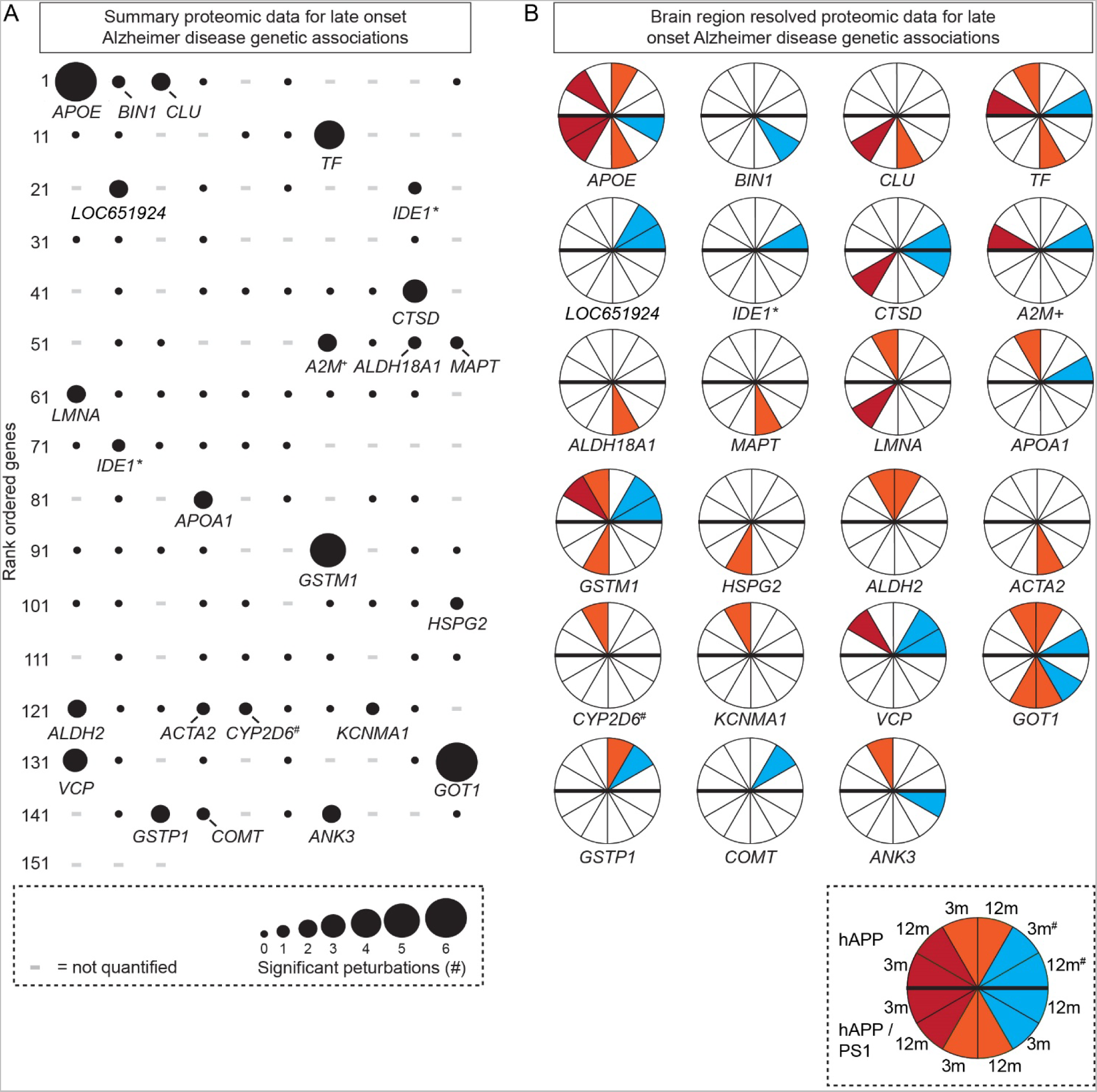
Assessment of Changes in Protein Abundance for Corresponding Genes Linked to Sporadic AD **(A)** Proteins with significant changes mapped to 153 genes previously identified as having a genetic association in human patients with sporadic AD. Size of the black circle shows the number of analysis paradigms that were identified as significant (FDR-adjusted *p* value < 0.05). - = not quantified. **(B)** Illustration of the specific brain region(s) and time point(s) that the significant changes were detected. Note: APOE was the top hit in the proteomics and was only identified as significantly altered in the Tg-AD mouse brain regions with increased Aβ levels. Red is for FC, orange for HIP, and teal for CB. # = hAPP cerebellum which lacks Aβ accumulation. * = IDE1* is shown twice because of multiple sequence variations, + = A2M+ was not identified but closely related alpha-2-macroglobulin protein Pzp is reported, # = CYP2D6# is not present in the mouse genome but we measured closely related Cyp2d22.

To focus our attention on the most confidently altered proteins in our large datasets and compare them between the two models, we generated rank ordered summary “confidence plots” for all the proteins found significantly altered (FDR-adjusted *p* value < 0.05). In the hAPP model, α-spectrin 2 (Spna2), which is known to have altered levels in AD, was the only protein identified as significantly altered in all of the four hAPP data sets with elevated Aβ (Figure 3A). We found 27 significantly altered proteins in three out of four hAPP datasets, 125 in two, and 691 in one. Similarly, only microtubule associated protein 2 (Map2) was identified as significantly altered in all six hAPP / PS1 data sets, and we found three in five, 20 in four, 36 in three, 214 in two, and 469 in one (Figure S2G). Next we compared the top 30 proteins (~3.5 - 4%) between the models which showed that 9 proteins met this criterion in both models. These mostly-confidently altered proteins included ApoE, Glial fibrillary acidic protein (GFAP), Solute Carrier Family 12 Member 5 (Slc12a5), neurofilament medium peptide (Nefm), clatherin heavy chain 1 (Cltc), Ankyrin 2 (Ank2), spectrin β-III (Spnb3), aconitase 2 (Aco2), and dynein cytoplasmic 1 heavy chain 1 (Dync1h1). These results are consistent with previous findings that the structural proteins Nefm and Spnb3 have altered levels in AD brains (Ciavardelli et al., 2010; Masliah et al., 1990). Slc12a5, which is the major extruder of chloride in neurons, plays a key role in shifting the effect of GABA from the depolarizing to the hyperpolarizing, and has been linked to epilepsy which may be relevant in AD (Lagostena et al., 2010; Palop and Mucke, 2009). Further, Ank2, GFAP, Aco2, Cnp, and Dync1h1 all have also been reported as altered in AD (Burbaeva et al., 2005; Kamphuis et al., 2014; Lazarov et al., 2002; Mangialasche et al., 2015; Silva et al., 2013; Soler-Lopez et al., 2011; Wang et al., 2003a).

**Figure 3.**
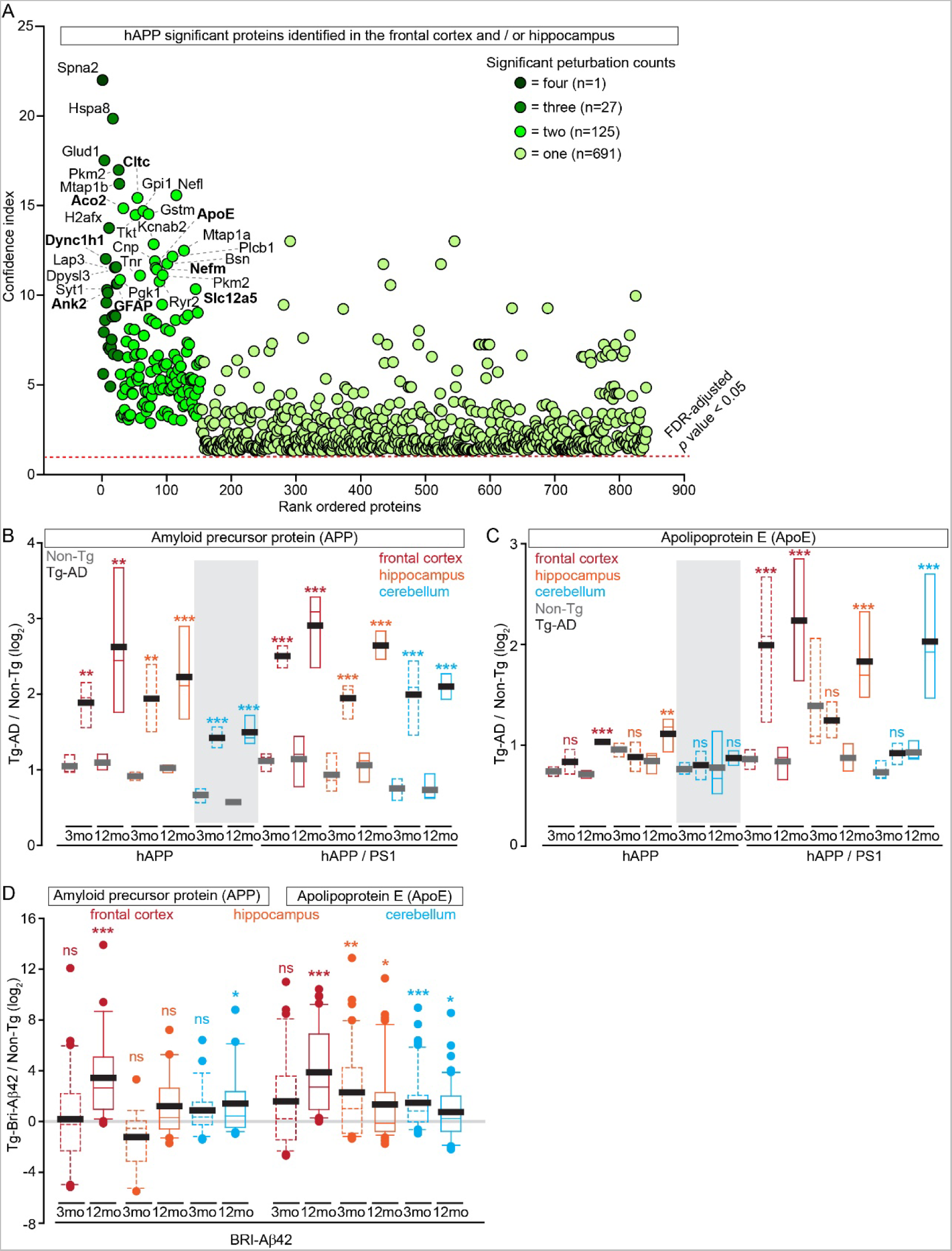
(See also Figures S2-3). Most Significantly Altered Proteins from the Brain Regions with Elevated Aβ Levels **(A)** Summary plot depicting the confidence index for all significantly altered proteins in the hAPP model in the four (pre- and post-symptomatic time points in the FC and HIP) brain regions with significantly increased Aβ levels. The number of datasets which each protein was identified in indicated by the shade of green, FDR-adjusted *p* value < 0.05. Proteins with very small *p* values > 0 but < 3.42 E-09 were graphed as 1E-13 (see Tables S1), proteins found in the top 30 proteins in both models are indicated in bold. **(B)** Analysis of APP abundance in Tg-AD datasets revealed a significant increase compared to Non-Tg controls. Magnitude of APP accumulation increased in each brain region from 3 to 12 mo. Only peptides with 100% sequence homology between mouse and human were included, n = 8 − 32 quantified peptides. **(C)** Apo E levels in hAPP were found significantly increased at 12 mo in FC and HIP but not CB. In hAPP / PS1 Apo E levels are significantly increased at 3 mo in the FC, and at 12 mo in all three brain regions. n = 17 − 39 quantified peptides. **(D)** APP and ApoE box plots showing protein levels from BRI-Aβ MS label free quantification at the indicated time points and brain regions. For (A-C) n = 3 mice per genotype at each time point and for (D) n = 4 mice, box plots defined by 25th percentile and 75th percentile. Median value is colored and mean is in grey or black, grey background indicates datasets without significantly increased Aβ levels. \**p* value < 0.05, \*\**p* value < 0.01, and \*\*\**p* value < 0.001 by Student’s t test for (B-C) and FDR-adjusted for (D).

Our finding that ApoE was among the most confidently altered proteins in our analysis motivated us to closely examine its expression patterns relative to APP and Aβ levels. Consistent with previous reports, in 3 mo hAPP mice we found on average that APP levels were increased ~2 fold in all brain regions and further elevated at 12 mo of age (Figure 3B). We obtained similar results for the hAPP / PS1 model and in general APP levels were slightly higher at both time points and in all brain regions compared to hAPP. Next we examined ApoE which in AD mouse models has been shown to be required for various Aβ pathologies (Bales et al., 1997). Interestingly, our analysis found that levels of ApoE were significantly increased at the post-symptomatic 12 mo time point in the FC and HIP of the hAPP model, and in the hAPP / PS1 model, ApoE had significantly increased levels at both time points in the FC and at the 12 mo time point in the HIP and CER (Figure 3C). Interestingly, ApoE levels were increased exclusively in those brain regions with significantly elevated Aβ levels, since it was unchanged at both time points in CER of hAPP mice which harbors increased APP abundance without an increase in Aβ levels (Figure S1A-B and Figure 3B). The positive correlation between high levels of Aβ and ApoE was further supported by our results showing that ApoE levels were increased in five out of the five possible post-symptomatic datasets with high Aβ levels.

To further explore this correlation and Aβ42 dependent proteome remodeling, we performed parallel label-free quantitative proteomic analysis of FC, HIP, and CER extracts at 3 and 12 mo time points of the BRI-Aβ42 AD mouse model (Figure S3A) (McGowan et al., 2005; Park et al., 2008). We hypothesized mice which over expresses Aβ42 fused to BRI transmembrane protein that require proteolytic cleavage by furin extracellular proteases to release Aβ42 may provide additional insight into Aβ dependent proteome remodeling since this model does not over express additional APP fragments. Further, these experiments should provide independent confirmation of our findings from the hAPP and hAPP /PS1 models. Consistent with previous reports, BRI-Aβ42 was only detected in mice carrying the transgene and overall levels increased over time (Figure S3B). In our proteomic analysis of BRI-Aβ42, we quantified more than 4,000 proteins in each dataset and identified 100’s of proteins with significantly altered levels (Figure S3C and Table S4). First we examined APP levels in the BRI-Aβ42 mice and found that in the majority of our datasets APP was relative unchanged save for the 12 mo time point in the FC and CER where we found a significant increase relative non-Tg littermates (Figure 3D). In contrast, we found ApoE levels to be consistently increased in all six BRI-Aβ42 datasets and significantly increased at 3 mo in the HIP and CER, and at 12 mo in all three brain regions (Figure 3D). Altogether, these results strongly suggest that Aβ42 accumulation leads to a concomitant increase in ApoE *in vivo* which likely has unexpected and important pathological consequences.

To test the confidence of our MS based protein abundance measurements we performed semi-quantitative western blot (WB) analyses of six selected candidate proteins. Indeed, these WB analyses unanimously confirmed proteins with both increased and decreased abundances (Figure S3D-H). Moreover, the BRI-Aβ42 proteomic analysis also provides additional independent confirmation for many proteins found altered in the hAPP and hAPP / PS1 models. Altogether, these results show that our proteomic analysis possesses the necessary analytical power and specificity to confidently reveal protein changes relevant to AD pathology.

### Age-dependent protein co-expression network analysis of Tg-AD brain region sub-proteomes

We hypothesized that consensus weighted gene / protein co-expression analysis (WGCNA), that identifies correlated patterns of protein levels in individual brain regions, across the two time points, and models would allow us to delineate modules (ME) of proteins that are potentially co-regulated in AD-like pathology (Langfelder and Horvath, 2007; Seyfried et al., 2017). We were able to successfully generate consensus weighted gene co-expression networks across two genotypes and time points for both the FC and HIP datasets but not for the CB. We identified 25 MEs for the FC and 11 MEs for the HIP across a total of 30 FC and 35 HIP datasets (Figures 4A, S4A, and Table S5). The FC MEs ranged in size from 25 to 842 proteins and the HIP MEs ranged from 26 to 900 proteins. In this way, WGCNA could reduce thousands of proteins measurements into 36 coherent protein MEs that represent core proteome remodeling programs that respond to an increasing Aβ load (Zhang et al., 2013). Direct comparison of the two topological overlap matrices from the Tg-AD or Non-Tg datasets showed that Aβ accumulation significantly (Z statistics *p* value < 1E-04), remodels several distinct molecular interaction networks in both the FC and HIP datasets (Figures 4B and S4B). To test if the MEs were enriched for proteins belonging to specific and previously described categories (Zhang et al., 2013), we subjected each ME to Gene Ontology cell component (GO:CC) enrichment analysis in R. Indeed, we identified significant gene ontologies assignments for 8 out of 11 HIP MEs which included myelin sheath (adjusted *p* value = 3.3E-08), regulation of actin polymerization (adjusted *p* value = 2.5E-07), mitochondrion (adjusted *p* value = 1.9E-17), synapse (adjusted *p* value = 4.09E-05), cell substrate adhereins junction (adjusted *p* value = 8.08E-05), and basal laminae (adjusted *p* value = 6.41E-04) (Table S5), which are consistent with previous findings in human AD (Bartzokis et al., 2007; Terry et al., 1991; Yamaguchi et al., 1992).

**Figure 4.**
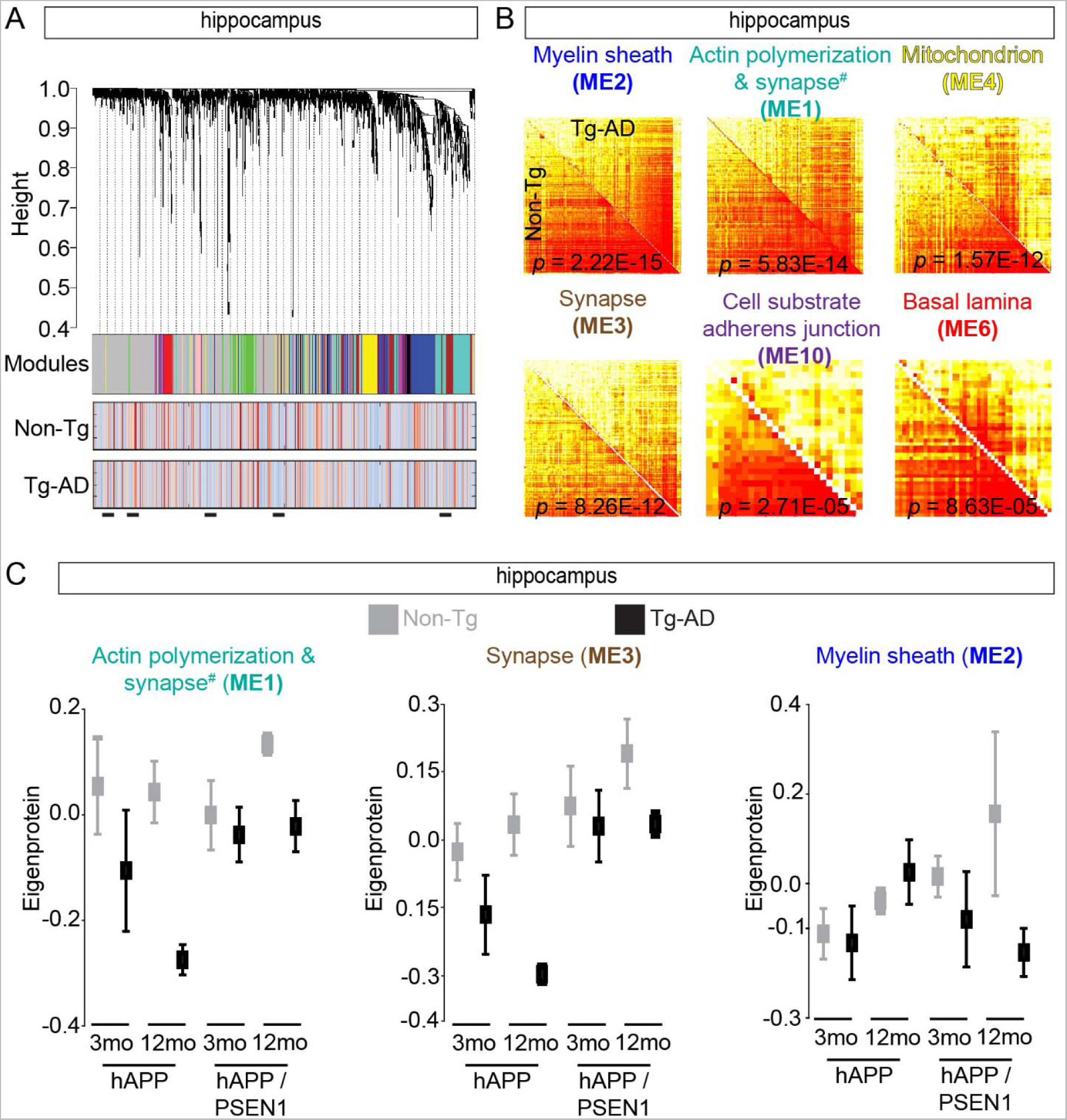
(See also Figures S4). Consensus Protein Coexpression Network Analysis of Tg-AD Models **(A)** HIP protein clustering trees from pooled data from both Tg-AD models and time points. Each module (cluster) is indicated by a distinct non-grey color just below the tree and grey represents proteins unassigned to an ME. The two rows of heat maps below show the association of individual proteins with *Q* for non-Tg or Tg-AD groups. Blue and red shading indicates proteins with reduced or increased expression, respectively, with increasing *Q*. Black bars indicate protein groups with differential expression. **(B)** Individual topological overlap matrices of significantly (Z statistics *p* value < 1E-04) differentially connected modules in HIP (C) between Tg-AD and Non-Tg datasets in both models. Module assignment based on the most significant GO assignment (^#^ for ME1 see Table S5). Shown are all modules with significant *p* values (< 1 E-4). **(C)** Summary expression value (eigenprotein) from the indicated HIP datasets, all Non-Tg / Tg-AD comparisons have *p* value < 0.05 calculated from Z statistics. Non-Tg is in grey and Tg-AD in black.

By decomposing our WGCNA modules into individual time points, we were able to identify shared and unique protein expression patterns between the two models (Zhang and Horvath, 2005), and by using a meta-analysis of correlations of ME eigenproteins (summary expression profile) (Langfelder et al., 2016), we were able to measure the relationship between a ME and each model at two time points. We found that six Hip and two FC MEs had significant (Z statistic and corresponding meta-analysis *p* value < 0.05), differences in abundance for each time point and model (Figures 4C and S4C-D). Consistent with previous results, we found both synapse related MEs (ME1 and ME3) in the HIP datasets to have reduced protein levels across all time points and models (Sheng et al., 2012). However, the myelin sheath (ME2) had only slightly altered expression patterns except at 12 mo in hAPP which showed a significant increase (Figure 4C). Parallel bioinformatic analysis of the FC MEs resulted in significant, but less dramatic eigenproteins (Figures S4D). Interestingly, only myelin sheath and mitochondrion MEs were identified in both the FC and HIP analyses suggesting that these processes are robustly perturbed in multiple brain regions (Table S5). Our network analysis provides new discovery-based information at the level of groups of co-expressed proteins in multiple Tg-AD models and brain regions. The strongest associations were found in HIP datasets, suggesting that Aβ accumulation affects co-expression of entire MEs more strongly in this brain region compared to FC or CB. Noticeably, in hAPP mice, the synapse MEs (ME1 and ME3) were significantly decreased at 3 mo (Figure 4C), which is before Aβ plaques could be detected (Figure S1D and S1K), and consistent with previous reports that the early synaptic deficits are caused by soluble Aβ peptides (Hsia et al., 1999). In contrast, the hAPP / PS1 model synapse MEs were only slightly reduced at 3 mo (Figure 4C), but Aβ plagues were already formed (Figure S1E and S1L). The hippocampal myelin sheath ME (ME2) also provides a contrasting view of the pathology in the two models (Figure 4C and S4D). In hAPP there were only slight differences in eigenprotein levels at both ages while in hAPP / PS1 mice we observed a substantial decrease at 3 mo and an even more dramatic reduction at 12 mo. This suggests that breaking down the myelin sheath may be more closely linked to Aβ plaques since the hAPP / PS1 mice had observable plaques at both time points (Figure S1E, S1H, S1L, and S1O). In summary, meta-analysis statistics allowed us to rank and prioritize each ME based on proteome expression patterns in two different Tg-AD modules and highlighted related but distinct changes in the synaptic proteomes of the hippocampus and frontal cortex.

### Refining the functional assignment of affected modules to untangle AD-like pathology

Our interpretation of the affected protein pathway perturbations in Tg AD hippocampus and frontal cortex is dependent on the confident identification of MEs. In order to identify the most confident groups of proteins in our datasets we first considered the most significant hippocampal MEs. ME2 and ME4 were the among the most significant and highly populated modules and GO:CC analysis clearly showed that these modules were mostly enriched in proteins involved with the myelin sheath (adjusted *p* value = 2.22E-15; n = 213) and mitochondria respectively (adjusted *p* value = 1.57E-12; n = 117). Since these proteins are already known to be impaired in AD, this finding shows that our analysis can efficiently reveal protein pathways relevant to AD (Bartzokis, 2004; Hirai et al., 2001). Thus, we focused on synapse related ME1 and ME3 from the hippocampus, and used Kyoto Encyclopedia of Genes and Genomes (KEGG) knowledge base to test if our modules are significantly enriched in proteins with shared functional assignments which may aid the interpretation of our data (Kanehisa and Goto, 2000). Interestingly, for ME1, when we compared the GO enrichment FDR versus the number of proteins in each category we found that long-term potentiation (LTP) to be the most enriched functional assignment based on more than 20 proteins (Figure 5A). This result is consistent with many previous reports which have shown that an impairment in hippocampal LTP may represent the basis of impaired synaptic plasticity and memory impairments seen in AD patients and models (Chapman et al., 1999), while ME3 was most enriched for the KEGG category assigned to synaptic vesicle cycling. Consistent with these results, when we used the GO: biological processes (GO:BP) database to further examine ME1 and ME3, we determined that these modules were significantly enriched for synapse plasticity / actin organization and modulation of synaptic transmission / vesicle transport / development (von Mering et al., 2005) (Figure 5B). These results suggest that synaptic transmission may be altered in the hippocampus by multiple mechanisms in AD-like pathology.

**Figure 5.**
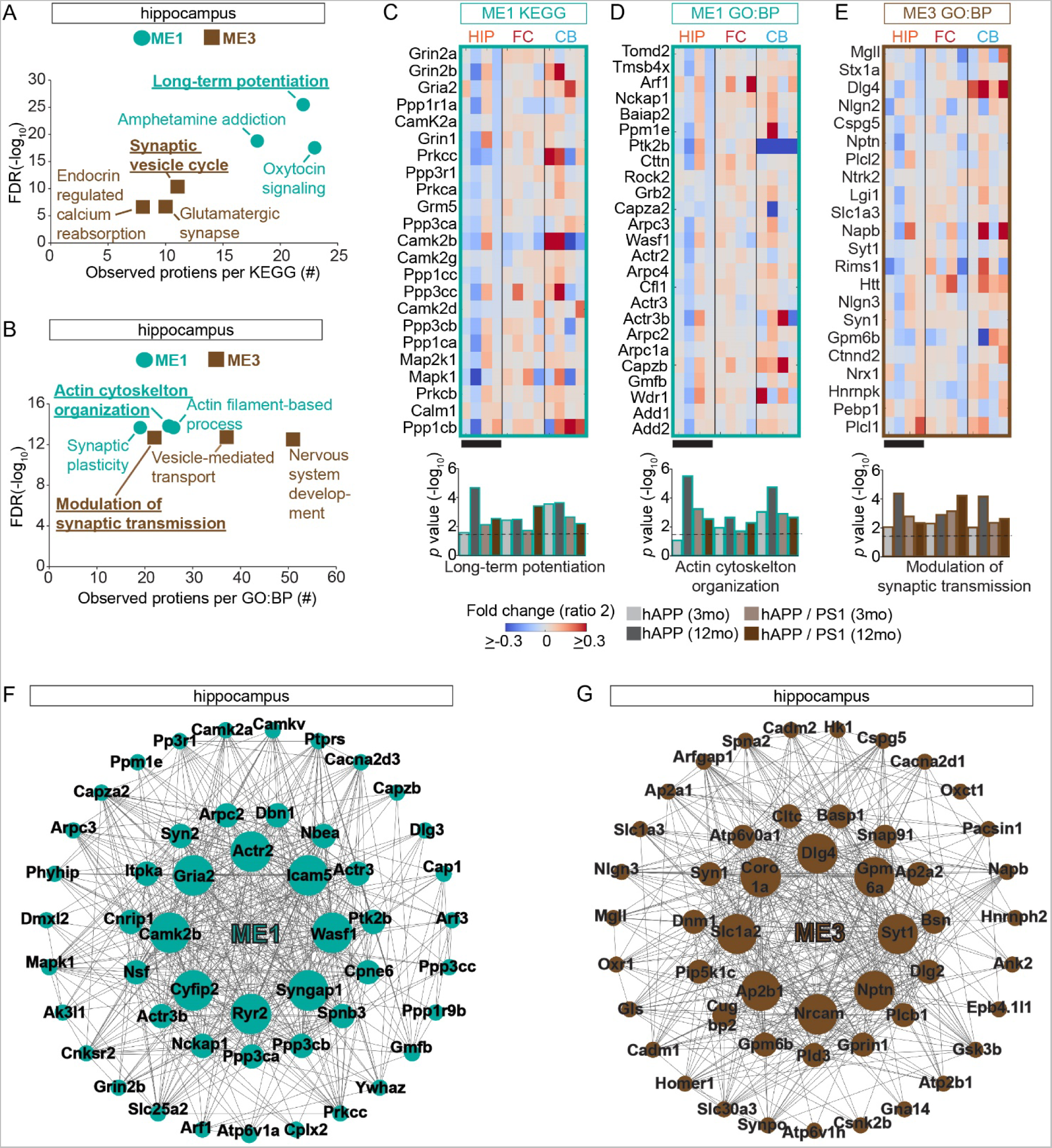
Synaptic Proteins in Module ME1 and ME3 Have Brain Region and Model Dependent Changes in Protein Levels **(A)** Hippocampal modules ME1 and ME3 are enriched in proteins which function in distinct synaptic functions based on KEGG. Shown is a scatter plot comparing the enrichment FDR (y–axis) versus the number of proteins per KEGG pathway (x-axis). The highly significant pathways are labeled and the most significant KEGG pathway for ME1 is long-term potentiation is underlined. **(B)** Hippocampal modules ME1 and ME3 are enriched in proteins which function in distinct synaptic functions based on GO : biological process (BP). As described for (A) except the BP, actin cytoskeleton organization was most significant for ME1 and modulation of synaptic transmission for ME3. **(C-E)** Protein expression matrix (ratio 2 see Figure 1) for the indicated modules for both Tg-AD models and time points. Black bars indicate the datasets with contrasting protein expression patterns. Below bar graph is based on the hypergeometric enrichment *p* values for each data set (brain region, model, and time point), and dotted line indicates an adjusted *p* value < 0.05. **(F-G)** Network of the top 50 hub proteins in ME 1 and ME3 from the hippocampus.

Next, to confirm the validity of this analysis strategy, we explored if any of our identified MEs relate to previously characterized AD pathology in the frontal cortex. We first used the KEGG database to investigate if any of our MEs are significantly enriched for the proteins in the KEGG AD pathway (hsa05010). Interestingly we found that ME1, ME2, and ME4 all are significantly enriched in proteins previously associated with AD based on KEGG. Then to extend these findings and further probe the potential functions of these MEs, we subjected these three MEs to GO:BP analysis. This analysis showed that these BP terms, namely ME1 - excitatory postsynaptic potential, ME2 – Oxidative-reduction process, and ME4 – hydrogen ion transporter activity were significantly enriched (Figure S4E). All of these pathways have been previously linked to AD pathology demonstrating that our data set has revealed relevant AD protein networks.

We also examined the identity and expression pattern of the individual proteins of the most significant MEs, which provided us with new insight into the core protein network perturbations occurring in AD-like pathology. First we inspected the hippocampal LTP module (ME1) which contained critical proteins such as Gria2, Grin2a/b, and Calm1, among others (Figure 5C). Overall, these proteins showed a progressive and significant reduction in expression levels in hAPP but were mostly up regulated at 3 mo and down regulated at 12 mo in hAPP / PS1. For the FC and CB in both models, proteins in this module generally had increased levels. We obtained similar results for the ME1 proteins associated with actin organization which is a process known to be important for synapse remodeling (Figure 5D). In general, the proteins comprising the second hippocampal synapse related module (ME3) showed a more robust down regulation of protein expression in hAPP compared to hAPP / PS1 which highlights potential differences in the synaptic pathology (Figure 5E). ME2 and ME4, which were identified in the FC data sets, are involved in mitochondrion based processes and were found to have mostly reduced expression in the HIP, but increased expression in the FC and CB datasets (Figures S4F-G). Finally, we performed network analysis based on the continuous measure of membership and connectivity based on WGCNA to determine the top 50 hub genes in ME 1 and ME3 from the hippocampus (Langfelder et al., 2016). These hub protein alterations are highly similar to the eigenproteins of these MEs and largely reflected biological function of corresponding MEs in AD. Interestingly, several of the core hub proteins identified in ME1 including Wasf1 and Nckap1 for example (Ceglia et al., 2015; Govek et al., 2005; Kim et al., 2006; Yamamoto et al., 2001), which, play critical roles regulating spine structure and have been implicated in AD (Figures 5F-G). Collectively, these analyses provide core protein modules of central importance to AD pathology.

### Validation of proteomic resource: AMPARs as core post-synaptic complexes altered in AD

Our WGCNA analysis identified the core AMPAR subunit Gria2 as a top hub gene of ME1 in the hippocampus in both mouse models. Additional Gria2 regulatory proteins such as Nsf and Camk2b (Braithwaite et al., 2002; Kristensen et al., 2011; Shanks et al., 2012), were also identified as top hub genes (Figure 5F). This finding strongly suggests that Gria2 containing protein networks may represent key potential targets for AD therapeutic intervention. However, Gria2 (even Nsf and Camk2b) play many essential synaptic functions in critical processes such as learning and memory and thus the possibility of directly manipulating their expression is too risky. Thus we focused on finding the key components of Gria2-centered meta-molecular complex contributing in AD. To explore this result and identify the earliest AMPAR alterations we examined the levels of all the AMPAR protein subunits in the HIP and FC datasets from hAPP mice at 3 and 12 mo. At 3 mo, the only significantly altered AMPAR proteins we identified were TARPγ-2 in the HIP dataset (1.253 + 0.162 versus 0.847 + 0.231 mean + STD, *p* value = 4.29 E-04) and TARPγ-3 in the FC (1.165 + 0.0354 versus 2.380 + 0.139 mean + STD, *p* value = 7.21 E-04) (Figure S5A). This result contrasts, our results at 12 mo, where we found both Gria1 and 2 significantly down regulated in the hippocampus, and we failed to quantitate either TARPγ-2 or 3 (Figure S5B). Based on the critically important effects that TARPs exert on AMPAR trafficking and synaptic responses, we next aimed to test if TARPγ-2 or 3 levels are also altered in the context of intact AMPARs in AD model brains (Tomita et al., 2005). Indeed, AMPARs immunopurified from hAPP brains with anti-Gria2 antibodies analyzed by WB and MS both showed decreased levels of TARPγ-2/3, strongly suggesting AMPARs in AD model brains have altered interactions with TARP proteins (Figures 6A and S5C). Finally, to test if AMPAR complexes are also remodeled in AD human brain tissue, we immunopurified AMPARs with anti-Gria2 antibodies and analyzed the purified protein complexes with MS and confirmed reduced TARPγ-2 levels in the context of human AD pathology (Figure 6B).

**Figure 6.**
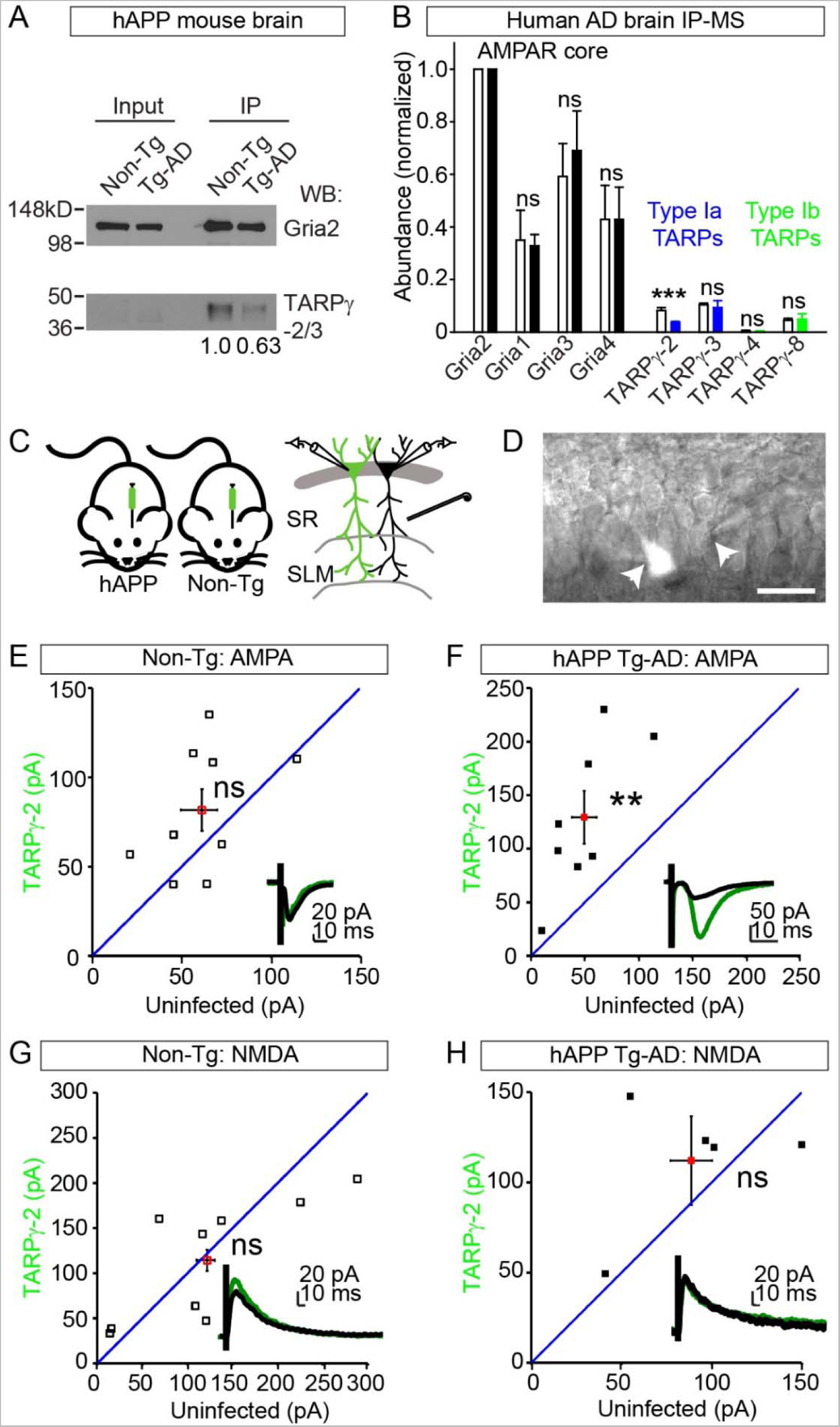
(See also Figures S5). AMPAR Complexes Represent Key Hub Proteins Altered Early in AD **(A)** Representative western blot analysis of AMAPR complexes immunoprecipitated from hAPP or non-TG mouse brain extracts with Gria2 antibodies. Less TARP-γ2 / 3 was recovered in the hAPP brain compared to Non-TG after normalizing the immunoprecipitated recovery of Gria2: 1.0 verses 0.63. **(B)** Semi-quantitative MS analysis of AMAPR complexes purified with Gria2 antibodies from human AD or healthy control brains show AMPARs in AD cortex have reduced levels of TARP-γ2 associated normalized to Gria2: healthy cortex = 0.082 + 0.011 verses AD cortex = 0.035 + 0.0058. Bar shows mean ± standard deviation, AD brains (n = 4 patients) and healthy controls (n = 2). White bars = healthy controls and solid bars = AD. **(C)** A lentivirus expressing Flag- TARP-γ2-IRES-GFP under the ubiquitin promoter was injected into CA1 in P1 mice. Whole cell recordings were made from neighboring infected and uninfected CA1 pyramidal cells in acute slices prepared from either hAPP mice or non-Tg littermates at P13-16. Cells were held at either -70 mV or +40 mV to measure AMPAR- and NMDAR-mediated currents, respectively in response to stimulation of the Schaffer Collaterals. **(D)** Example image of simultaneously recorded infected and uninfected CA1 pyramidal cells in a hAPP hippocampal brain slices. **(E)** AMPA currents are not significantly different in TARP-γ2-infected versus uninfected cells in non-Tg littermates (Mean ± SEM: Stg: 70.248 ± 11.479; Uninf: 59.896 ± 11.020, n=7, *p* value = 0.379). **(F)** AMPA currents are significantly increased in TARP-γ2 infected cells relative to uninfected neighboring cells in hAPP mice (Mean ± SEM Inf: 134.67 ± 27.812 pA; Uninf: 48.331 pA, n = 7, *p* value = 0.00385). **(G)** NMDA mediated currents are not significantly different in TARPγ−2 infected versus neighboring uninfected neurons in WT mice (TARPγ−2: 114.2344 ± 22.5515, Uninf: 121.669 ± 29.973, n = 9, *p* value = 0.7089). **(H)** NMDA mediated currents are not significantly different in TARPγ−2 infected versus neighboring uninfected cells in hAPP animals (Mean ± SEM: TARPγ−2: 112.148 ± 16.537; Uninf: 88.6866 ± 0.301, n = 5, *p* value = 0.3016). \*\**p* value < 0.01, and \*\*\**p* value < 0.001 by Student’s t test.

Impaired fast excitatory synaptic transmission and AMPAR dysfunction has previously been implicated in the AD hippocampus, however the details have remained unclear (Hsia et al., 1999; Hsieh et al., 2006). Consistent with previous observations, we also observed a significant reduction in the strength of evoked field potentials in hAPP hippocampal slices relative to control littermates (Figures S5D-F). We hypothesized that reduction of TARPγ-2 in AD could lead to destabilization of synaptic AMPARs since TARPγ-2 anchors AMPAR subunits to the first two PDZ domains of PSD-95, and the interaction between PSD-95 and TARPγ-2 can help deliver AMPARs to the synapse (Schnell et al., 2002). Although increasing the levels of TARPγ-2 increases the amount of extrasynaptic AMPARs without affecting basal AMPAR-mediated synaptic transmission in wild type cells (Schnell et al., 2002), we wondered if TARPγ-2 might affect basal synaptic transmission in hAPP mice. To test this, we generated lentiviruses that overexpress TARPγ-2 and cytosolic GFP in subsets of neurons and injected this virus into CA1 in transgenic and control littermates (Figures 6C-D and S5G). Consistent with previous observations (Schnell et al., 2002), we found that overexpression of TARPγ-2 had no effect on basal AMPAR EPSCs in non-Tg animals, but surprisingly, we found that TARPγ-2 overexpression resulted in a significant increase in AMPAR EPSCs in transgenic hAPP mice (Figures 6E-F). Additionally, we observed no effects on NMDAR EPSCs in either control or transgenic mice, consistent with TARPγ-2 selectively regulating AMPA-type glutamate receptors (Figures 6G-H). These results support TARPγ-2 as potentially safe target for targeting in AD since the effect on healthy synapses is limited. Our results highlight the potential importance of this AD proteomic resource by functionally validating a pioneering observation from our proteomic data that provides a new mechanism that may explain impaired excitatory synaptic transmission in AD.

## Discussion

Global quantitative proteomic analysis of brain tissue extracts from hAPP, hAPP / PS1, and BRI-Aβ transgenic mouse models of AD-like pathology have highlighted the complex molecular and cellular responses of altered APP processing and elevated levels of Aβ peptides on different brain regions *in vivo* and identified many altered proteins that have yet to be reported. Of the three brain regions analyzed, we found the hippocampus to possess the largest number of proteins with altered abundances. All three brain regions showed a progressive age dependent increase in proteome remodeling, and a small collection of proteins were found altered in multiple brain regions and in multiple models. Overall more proteins were found to have increased rather than reduced levels in Tg-AD brains, compared to littermate controls. The abundance of ApoE protein in AD and AD models have been uncertain, we found that ApoE to have increased levels only in those brain regions and models with high levels of Aβ (Gupta et al., 2011; Hesse et al., 2000; Kuo et al., 2000; Liu et al., 2013; Liu et al., 2007)Comprehensive protein co-expression analysis identified several distinct modules enriched for proteins involved with myelin, synapses, and mitochondrion. Additional bioinformatics analysis showed that two distinct synapse modules have different expression patterns which suggest that synapses may be altered through more than one mechanisms in AD. AMPAR subunit Gria2 was identified as a hub protein and biochemical purification of AMPARs from mouse and human AD brains show that auxiliary type 1a TARPs are perturbed early, while Gria1 and Gria2 become reduced later in disease progression. Correcting TARPγ-2 levels in young mice can correct AMPAR mediated synaptic transmission in hAPP mice.

### Toxic Aβ levels drive non-linear proteome remodeling

Our proteomic characterization of AD model brains showed that elevated Aβ levels cause major alterations of the brain proteome and significantly alters the level of many proteins expressed in multiple cell types that function in distinct pathways. Our results also show that the toxic effects of Aβ on the brain are extensive and non-linear. For example, if we just consider the most confidently altered proteins (Figure 3A and S2G), it is clear that multiple cell types and protein networks are affected by toxic levels of Aβ. For example, we identify several neuronally expressed proteins with altered levels such as the dendritic protein Map2 and synaptic protein Bsn which plays a key role in organizing the cytomatrix of a subset of glutamergic synapses which may be particularly important in AD (Altrock et al., 2003). Additionally, among the most confidently identified altered proteins, we also identified proteins that are expressed in astrocytes such as GFAP which responds to insult and injury in the CNS (Eng et al., 2000), and Hba-a1 which is expressed in neurons, glia, and oligodendrocytes (Biagioli et al., 2009). Our WGCNA analysis identified modules involved with synapses and myelin which also supports the idea that multiple cell types are altered by AD pathology (Figure 4 and 5). These results provide discovery-based support of the hypothesis that Aβ pathology is highly complex, non-linear, and broadly affects many proteins and cell types in the brain (De Strooper and Karran, 2016).

ApoE is predominantly produced by glia cells in the mammalian central nervous system and is believed to be involved in multiple mechanisms important to AD including Aβ clearance and aggregation, signaling, lipid metabolism, and synaptic plasticity (Holtzman et al., 2012). Our quantitative proteomic results from three distinct mouse models of AD-like pathology showed that ApoE levels are specifically increased in those brain regions that harbor increased levels of Aβ, while we failed to detect a significant increase in ApoE levels in the cerebellum of hAPP mice suggesting that it is primary involved in Aβ clearance rather than production (Figure 3C). This result contrasts previous results showing that the APP intracellular domain can drive ApoE gene transcription and may highlight differences between AD models and the complexity of ApoE biology (Liu et al., 2007). A variety of ApoE receptors localize to the post synaptic membrane and include Lrp1, Apoer2, and Vldlr which interact with several synaptic scaffolds and glutamate receptors (Lane-Donovan et al., 2016). Recently, an allele specific role of ApoE in regulating complement C1q protein has been reported and shown to play an important role in regulating the rate of synaptic pruning (Chung et al., 2016). It is possible that the synaptic protein defects which we identified could be due to altered ApoE levels at tripartite synapses.

### AMPARs as candidate examples for AD proteomic resource

There have been many previous reports describing impaired synaptic transmission and an overall reduction in the number of synapses in the hippocampus of AD models and patients long before the appearance of detectable plaques (Shankar et al., 2008; Terry et al., 1991). Consistently, we confirmed that distinct components of excitatory synaptic proteomes are significantly remodeled before plaque formation (Figure 4C). While, a universal and unifying understanding of precisely how synapses become altered in AD has been lacking, it is possible that Aβ impairs synaptic function through direct and indirect mechanisms (Sheng et al., 2012). Indeed, WGCNA analysis of the HIP from the hAPP and hAPP / PS1 models revealed two synaptic modules with reduced protein levels at early and late time points (Figures 4 and 5). ME1 is enriched in proteins involved with the processes of long-term potentiation and actin skeleton organization, both of which have been implicated as playing key roles in AD (Sheng et al., 2012). ME3 is enriched in proteins involved with the modulation of synaptic transmission (Figure 5A-B). Among the top 50 hub proteins in ME1 were AMPA and NMDA receptor complex subunits (Figure 5F), and close examination of AMPAR components showed that pioneering reduction in protein abundance was observed in the critical auxiliary subunit TARPγ- 2/3 and it was not until the later time point that Gria1 and 2 had significantly reduced levels in the hippocampus of hAPP mice (Figure S5A-B). It has recently been shown that AMPAR density was significantly reduced at perforated synapses and synapses onto parvalbumin-positive neurons in the CA1 region of TARPγ-2 KO mice which is interesting since we found significantly reduced levels of parvalbumin in the hippocampus of both models (Table S3) (Yamasaki et al., 2016). Furthermore, pore forming Gria3 and 4 AMPAR subunits were not found as being significantly altered, however in other AD models they have been showed to be important Aβ substrates (Reinders et al., 2016) (Figure S5A-B). AMPAR downscaling and removal have been previously described in multiple AD models and has been shown to occur before plaque formation but the precise mechanism explaining these observations have remained vague (Chang et al., 2006; Hsia et al., 1999; Hsieh et al., 2006). These findings motivated us to test whether restoring TARPγ-2 levels could rescue the previously reported AMPA defect in hAPP mice. Indeed, we found that over expression of TARPγ-2 caused a large increase in AMPAR- but not NMDAR-mediated currents, which functionally validates a top candidate protein identified in our analysis (Figures 6E-H). Altogether, our results suggest that Gria1, 2 containing AMPARs which are known to be enriched in TARPγ-2 and 3 may represent key Aβ targets due to their restricted expression patterns at the effected synapses in the hippocampus (Schwenk et al., 2012). Finally, while both NMDA and AMPA receptors play critical roles in the establishment and maintenance of LTP, our results highlight a new potential mechanism by which reduced expression of TARPγ-2 would result in impaired delivery of AMPARs to spines and thus compromise LTP in the hippocampus of AD patients. TARP subtypes have been shown to differentially influence AMPAR gating (Milstein et al., 2007), and TARPγ-2 may specifically represent a therapeutic target to restore cognitive function in AD patients.

### Temporal-spatial resolved atlas of proteome remodeling in mouse models of AD-like pathology

We set out to determine the protein substrates and mechanisms of Aβ toxicity in brain and to dissect the amyloid cascade hypothesis by using three distinct and complementary mouse models of AD-like pathology. Our large-scale proteomic analysis shows that Aβ alters multiple protein networks associated with synapses, mitochondrion, and myelin through direct and indirect mechanisms. In order to maximize the accessibility and impact of our large scale proteomic results, we have generated a new online interactive AD model protein expression database (PINT, Proteomics INTegrator) as a resource for the entire AD research community http://sealion.scripps.edu/pint/?project=3d7c1ac078930a798a07c6a397bd21ef. This resource allows users to query proteins of interest, visualize the quantified peptides, and perform enrichment analyses (Figure S6). While several previous studies have reported global analysis of mRNA levels from AD model and patient brains, here we report a more directly relevant measure of the hampered biological processes in brain. Moreover, by determining the abundance of thousands of proteins from three mouse models of AD-like pathology at pre- and post-symptomatic stages, we revealed many new temporal and spatially restricted protein alterations. The importance of determining protein abundances rather than mRNA levels is particularly relevant for the investigation of AD since it is characterized by an altered proteostasis network and reduced protein degradation capacities (Powers et al., 2009; Rubinsztein, 2006). However, our study is not without limitations since we failed to accurately quantify many proteins expressed at low levels. Unfortunately, our results are also limited by protein abundance averaging between the multiple cell types present in the brain and we can only draw conclusions for those proteins we found significantly altered.

In summary, our results show that Aβ accumulation causes a broad and progressive alteration in the abundance of many functionally and spatially linked components of the brain proteome. Furthermore these findings and those recently published from others (Seyfried et al., 2017), raise the possibility that studying groups of co-expressed proteins, or modules, might have a strategic advantage over the study of individual proteins due to the highly complex response of various cell types to toxic levels of Aβ.

## Experimental Procedures

See Supplemental Information

## Author Contributions

J.N.S, Y.Z.W., D.B.m.C., L.A.D., E.H.K., T.N., A.G., and J.R.Y. designed research; J.N.S., Y.Z.W., S.M.B., L.A.D., D.B.m.C., N.F.S., T.J.H., K.A.C., S.N.S., and M.L.A. performed research; J.N.S., Y.Z.W., S.M.B., S.K.P., M.L.A., E.H.K., E.M., and J.R.Y. contributed new reagents/analytical tools; J.N.S, Y.Z.W., L.A.D., S.M.B., N.F.S., M.L.A., T.J.H., K.A.C., S.K.P., J.W.K., T.N., A.G., and J.R.Y. analyzed data; J.N.S. and J.R.Y. wrote the paper.

## Acknowledgments

We thank Lars Bertram, Floyd Sarsoza, Ianessa Morantte, Erik Kapernick, Joris De Wit, Lujian Liao, Claire Delahunty, Tao Xu, Edward Rockenstein, Jaehyuk Choi, Andrew Dillin and Chandra Inglis for their assistance on this project. Imaging was performed at the Northwestern University Center for Advanced Microscopy generously supported by NCI CCSG P30 CA060553 awarded to the Robert H Lurie Comprehensive Cancer Center. Histology services were provided by the Northwestern University Mouse Histology and Phenotyping Laboratory which is supported by NCI P30-CA060553 awarded to the Robert H Lurie Comprehensive Cancer Center. This work was supported by NIH awards F32AG039127 and R00DC013805-02 (J.N.S.); R01 MH067880, P41 GM103533 (JRY). M.L.A. held a postdoctoral fellowship from Fonds de Recherche du Québec—Nature et Technologies. Upon acceptance of the paper the raw MS files will be accessible though the Massive repository and ProteomeXchange consortium (Accession: PXD005595, FTP Download: ftp://MSV000080431@massive.ucsd.edu/ http://proteomecentral.proteomexchange.org/cgi/GetDataset?ID=PXD005595).

